# Dynamic allocation of orthogonal ribosomes facilitates uncoupling of co-expressed genes

**DOI:** 10.1101/138362

**Authors:** Alexander P.S. Darlington, Juhyun Kim, José I. Jiménez, Declan G. Bates

## Abstract

Introduction of synthetic circuits into host microbes creates competition between circuit and host genes for shared cellular resources, such as RNA polymerases and ribosomes. This can lead to the emergence of unwanted coupling between the expression of different genes, complicating circuit design and potentially leading to circuit failure. Here we demonstrate the ability of orthogonal ribosomes to alleviate the effects of this resource competition. We partition the ribosome pool by expressing an engineered 16S RNA with altered specificity, and use this division of specificity to build simple resource allocators which reduce the level of ribosome-mediated gene coupling. We then design and implement a dynamic resource allocation controller, which acts to increase orthogonal ribosome production as the demand for translational resources by a synthetic circuit increases. Our results highlight the potential of dynamic translational resource allocation as a means of minimising the impact of cellular limitations on the function of synthetic circuitry.

## Introduction

A key goal of synthetic biology is the construction of novel information processing genetic circuits in microbes which can be used to guide cellular function and control metabolic processes [1]. However, many initial designs fail upon implementation *in vivo* due to unforeseen interactions between the host and synthetic circuit [2]. These often arise due to competition for shared cellular resources, such as RNA polymerases and ribosomes [3]. Previous experimental studies have shown that translational capacity, in the form of free ribosomes, limits microbial gene expression [4, 5, 6, 7, 8] and so is a key cause of these hidden interactions [3]. Previous work has shown that the *non regulatory* coupling which emerges due to these hidden interactions can be reduced by careful design and selection of ribosome binding sites and plasmid copy numbers [3, 5, 6]. Additionally, incorporating negative feedback loops into the circuit can insulate genes [9, 10, 11]. These approaches, however, require significant re-design of the original synthetic circuit, and by incorporating additional regulatory interactions may make certain circuit behaviours unobtainable. In this work, we propose an alternative approach, based on the partitioning of the cell’s translational capacity, and show that it allows the decoupling of circuit genes from one another and from their host without the need for extensive modification.

Previously, both transcription and protein degradation activities have been partitioned into circuit-specific and host-specific activities through the use of ‘orthogonal’ components. For example, the expression of RNA polymerases (RNAP) from bacteriophage T7 in *Escherichia coli* creates a circuit-specific transcription system [12]. Co-option of proteases from other bacteria has been used to create a circuit-specific degradation pathway [13]. Due to the universality and complexity of the cell’s translational machinery, there does not exist a sufficiently distinct ribosome which can be co-opted into *E. coli* to create a truly orthogonal ribosome pool. However, translational capacity can be divided into host and circuit specific functions by the use of synthetic ribosomal components to create a quasi-orthogonal ribosome (‘o-ribosomes’) system [14, 15, 16]. The binding interactions between an mRNA and the 16S rRNA of the small ribosomal subunit are known to be a key regulator of translation initiation [17], and thus o-ribosomes can be created by expressing a synthetic 16S rRNA. Evolving or designing the 5’ sequences at and around the ribosome binding site (RBS) of circuit mRNAs to interact with this synthetic sequence allows the creation of an orthogonal translation system [15, 16]. For simplicity, we refer to this synthetic 5’ sequence as an orthogonal RBS (‘o-RBS’). These specialised o-ribosomes have previously been successfully used to probe various aspects of ribosome function [18] but their use in the creation of synthetic gene circuits has not yet been explored.

## Results

### Development of an o-ribosome model

We initially developed a mathematical model to assess the feasibility of implementing an orthogonal translation system, based on the microbial physiology model developed by Weiße *et al.* [19]. This model captures the three fundamental trade-offs in bacterial gene expression: (i) energy production is limited by substrate import and enzymatic activity, (ii) finite translational capacity, and (iii) the proteome mass is finite. We refined the original ribosome synthesis description by considering the separate production of a protein component, which we term the ‘empty ribosome’, and an rRNA component. The protein component is encoded by an mRNA and translated by host ribosomes. This species represents the small and large ribosomal subunits and other accessory protein complexes. The host and orthogonal rRNAs are transcribed and bind with the empty ribosome to form either functional host or orthogonal ribosomes. See Section S3.1 for the full model derivation. Our model predicts that cells can tolerate the use of orthogonal ribosomes for significant levels of gene expression (Figure S21a,b) and that partitioning of the ribosome pool can be used to divert translational machinery to a synthetic gene (Figure S21c).

### Construction of an orthogonal gene expression system *in vivo*

We utilised a previously described o-16S rRNA system, which contains an o-16S rRNA under the control of *P_lac_*, thus allowing its levels to be controlled by isopropyl *β*-D-1-thiogalactopyranoside (IPTG) [20]. Our circuit is carried on a second plasmid and consists of RFP under the control of the *P_lux_* promoter. Translation by either the host (h-RFP) or orthogonal (o-RFP) ribosome pools is controlled by selection of the RBS (Figure 1a). RFP mRNA production is induced with N-acyl homoserine lactone (AHL) via luxR which is constitutively expressed from the circuit plasmid and utilises host ribosomes for its expression.

**Figure 1:**
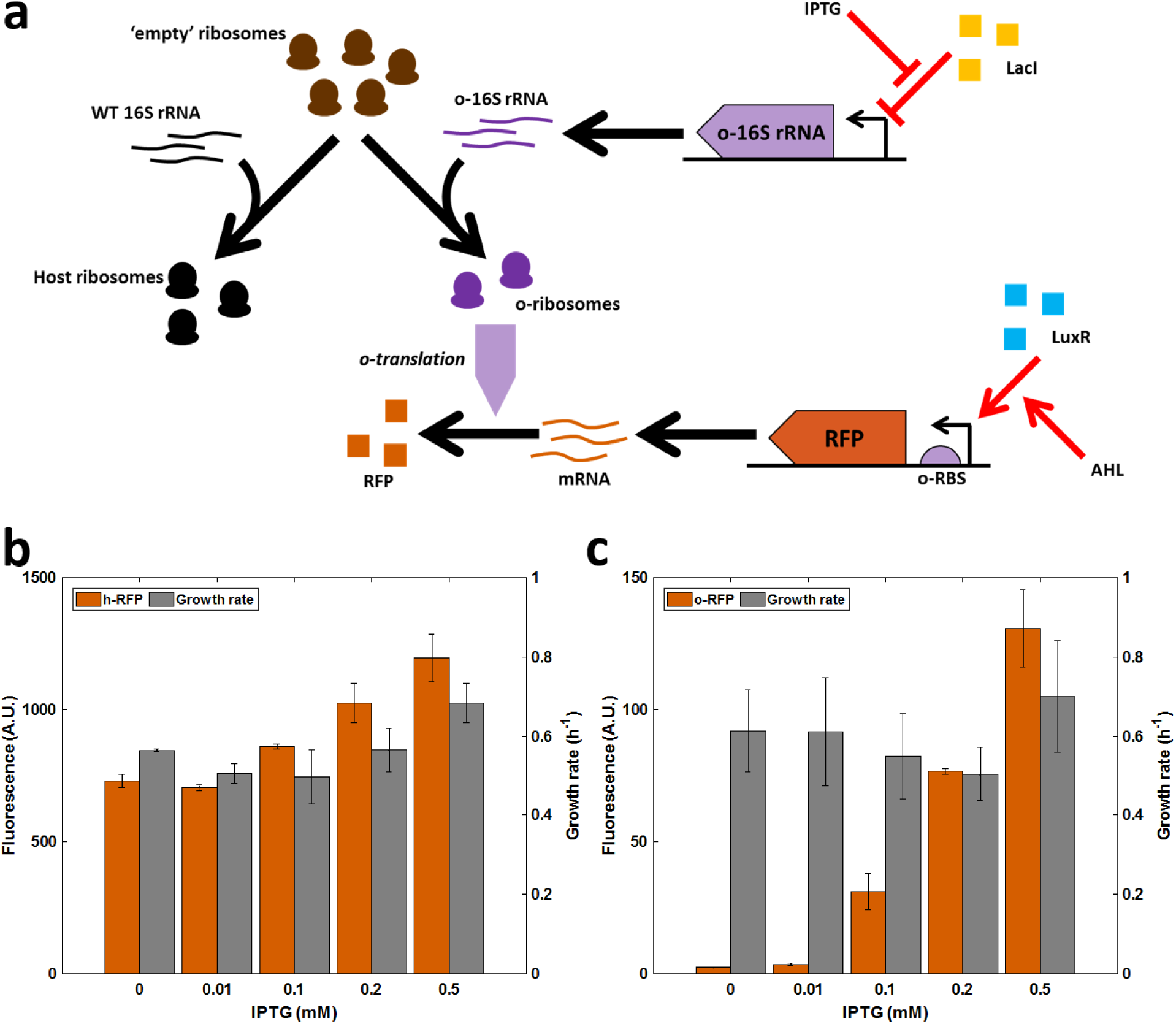
Developing the o-ribosome pool as an expression system. RFP expression using either the host or o-ribosome pool. o-ribosomes are produced by inducing o-16S rRNA with increasing concentrations of IPTG. (o/h)-RFP is induced with 20 nM AHL throughout. Bars represent means ± 1 SD. **a.** Schematic of the orthogonal translation system. **b.** Expression of h-RFP by the host ribosome pool. **c.** Expression of o-RFP by the o-ribosome pool.

To assess the impact of o-ribosome production on the host growth rate and gene expression, gene induction was maintained using a constant concentration of AHL in the presence of increasing IPTG concentration. The production of o-ribosomes alone has no effect on growth rate demonstrating that their presence is non-toxic (Figure S2). We observe a 50% increase in h-RFP fluorescence (*p* < 0.05, t-test, 0 mM *vs.* 0.5 mM IPTG) (Figure 1b). This is likely due to a low affinity of the o-ribosome pool for the host RBS which gives rise to a low level of ‘crosstalk’. Increasing the ability of genes to sequester ribosomes, for example by increasing RBS strength or expression level, abolishes the impact of cross talk (Section S3.5). Increasing the size of the o-ribosome pool acts to dramatically increase o-RFP expression, demonstrating the utility of o-ribosomes to set a ‘circuit-specific protein budget’ (Figure 1c). Use of o-ribosomes for gene expression has no effect on exponential growth rate (*p* = *N S* for pairwise comparisons at all IPTG levels). Expression of o-RFP is on the order of tenfold smaller than that of the h-RFP. This may be due to inefficient o-ribosome assembly resulting in a smaller number of available ribosomes or due to the difference in strengths of the o-RBS (Figure S23).

### Gene coupling in circuits utilising an orthogonal ribosome pool

Here we consider a new circuit consisting of two genes; the original RFP cassette and a new GFP cassette. GFP transcription was constitutively driven from the *P_tet_* promoter and the host or o-ribosome pool utilised for translation, controlled by selection of RBS as described above. We determined the level of coupling between the two circuit genes by observing the slope of the isocost line of circuit gene expression during exponential growth; this quantifies the change in GFP as RFP is induced [6] (Figure 2). During balanced exponential growth, the concentration of RNAP and ribosomes is constant, creating a ‘fixed protein budget’. This is shared across the circuit genes so that as more is ‘spent’ on one gene, less can be ‘spent’ on another. This interaction is quantified in the slope of the isocost line, [6], which demonstrates the potential combinations in which the two proteins can be produced given the fixed budget. Gene coupling in the h-RFP-hGFP circuit (utilising the host ribosome pool) results in a slope of −3.3; for every unit of RFP gained, ∼3 units of GFP are lost (Figure 2b). Tuning the o-ribosome pool when utilising the host ribosomes as the translational resource has no effect on the coupling observed, consistent with our model predictions (Figure 2a). Replacing the host RBS sequences with the o-RBS to produce the o-RFP-o-GFP circuit results in the coupling being reduced to 30% of that observed when using the host pool (Figure 2d). Increasing IPTG increases gene expression but has negligible effect on coupling, as predicted by our model (Figure 2b).

**Figure 2:**
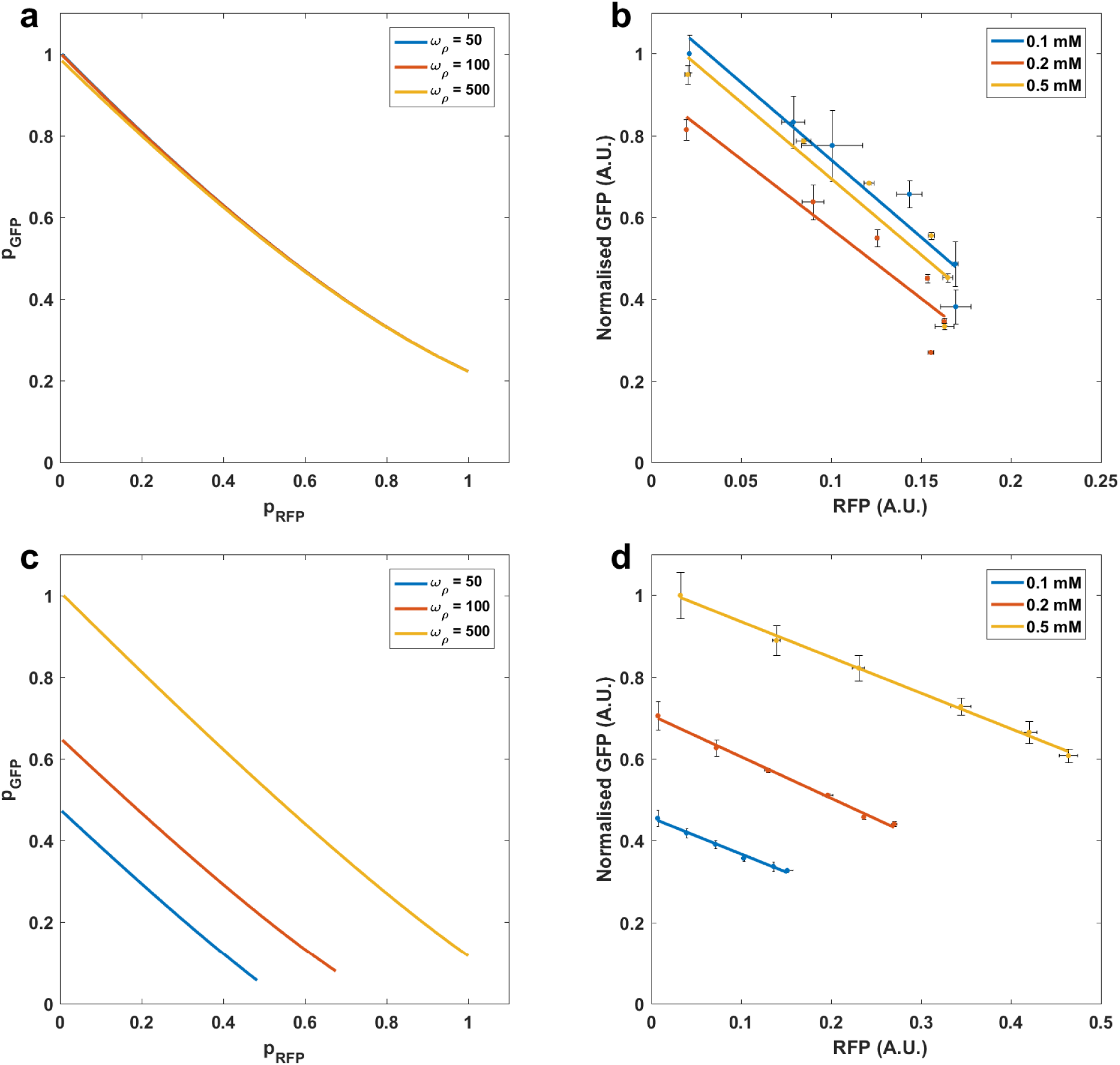
Gene coupling in gene circuits utilising either the host or orthogonal ribosome pool. Simulations of the steady state concentrations of RFP and GFP normalised by the maximum protein production achieved across the o-rRNA transcription rates tested. *ω_GF P_*= 100 and *ω_RF P_* = 1 to 10^3^ mRNAs per min. o-rRNA production (*ω_ρ)_* was simulated at the RNAs per min as shown. Experimental data was produced by inducing RFP using AHL from 0 to 20 nM. Points are the mean steady state fluorescence ± 1 S.D normalised by maximum GFP expression obtained across different levels of IPTG treatment. The isocost line is fit to the mean fluorescence. o-ribosome production was induced using IPTG at the concentrations as shown. **a.** Simulations of circuit using the host ribosome pool. **b.** *In vivo* protein productions using the host ribosome pool. **c.** Simulations of the circuit using the o-ribosome pool for translation. **d.** *In vivo* coupling observed in the circuit using the o-ribosome pool.

### Utilising both host and orthogonal ribosome pools reduces coupling of co-expressed genes

Having successfully partitioned the ribosome pool, we tested the ability of these pools to act as a simple distribution mechanism for translational capacity. Maintaining the original circuit topology and function, we altered the RBS of each gene to create two new circuit variants; o-RFP-h-GFP and h-RFP-o-GFP. Our model predicts that placing the constitutively expressed GFP under control of the o-ribosome pool (h-RFP-o-GFP arrangement) acts to insulate the gene from competition with RFP and so significantly reduces gene coupling, over a range of o-ribosome pool sizes (Figure 3a). Experimental validation of these predictions showed near complete abolition of the isocost line slope with coupling falling by over 95% (Figure 3b). Varying IPTG levels shows this decoupling is highly robust, with IPTG acting only to tune expression (Figure 3b).

**Figure 3:**
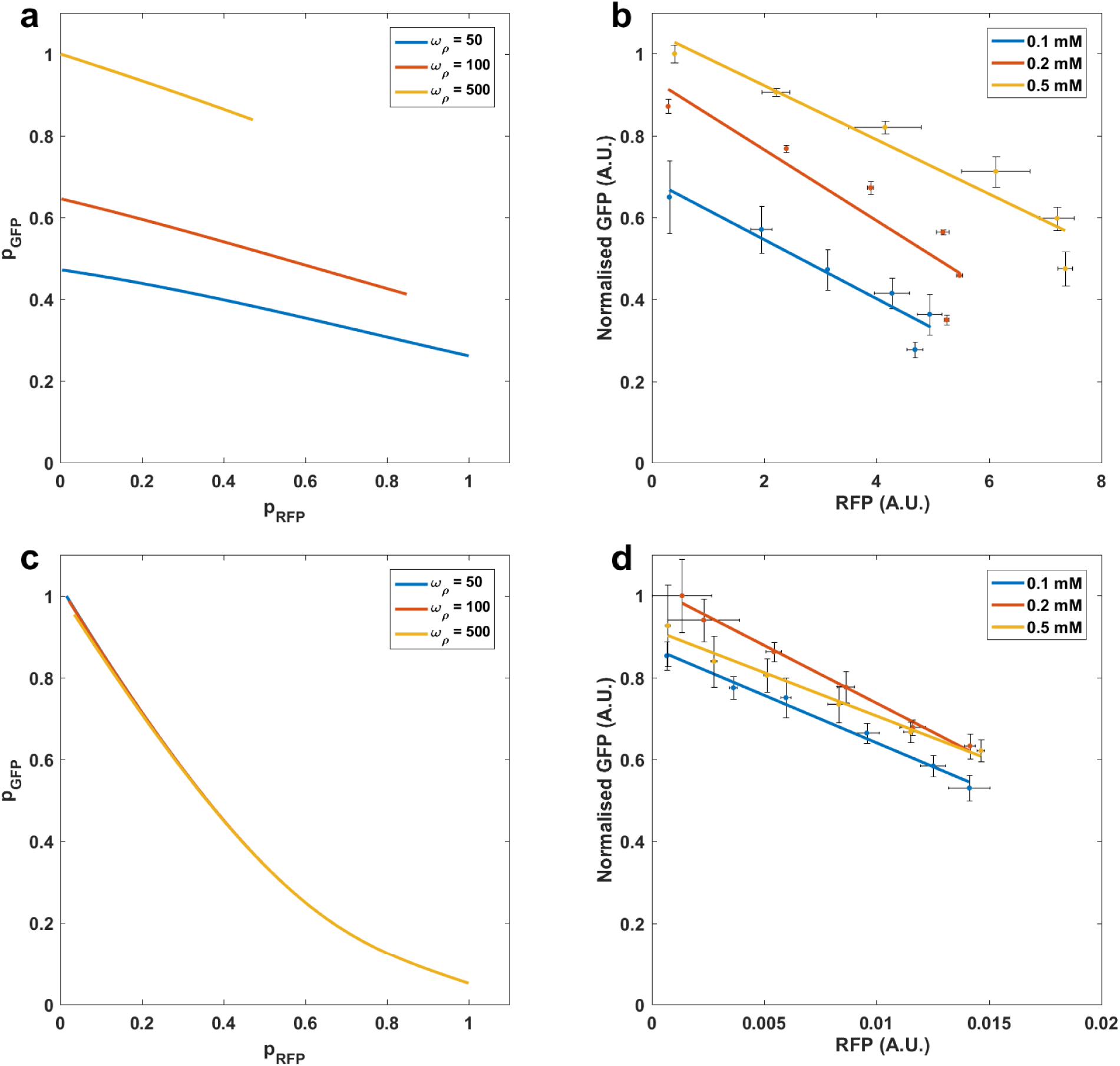
Gene coupling in circuits utilising different ribosome pools. Simulations of the steady state concentrations of RFP and GFP normalised by the maximum protein production achieved across the o-rRNA transcription rates tested. *ω_GF P_* = 100 and *ω_RFP_* = 1 to 10^3^ mRNAs per min. o-rRNA production (*ω_ρ_*) was simulated at the rates shown. Experimental data was produced by inducing RFP using AHL from 0 to 20 nM. Points are mean steady state fluorescence ± 1 S.D. normalised by maximum GFP expression obtained across different levels of IPTG treatment. The isocost line is fit to the mean fluorescence. o-ribosome production was induced using IPTG at the concentrations as shown. **a.** Simulations of the circuit using host ribosomes for RFP expression and o-ribosome for GFP expression. **b.** *In vivo* coupling of the h-RFP-o-GFP circuit. **c.** Simulations of the circuit using the o-ribosome for RFP expression and host pool for GFP. **d.** *In vivo* coupling of the o-RFP-h-GFP circuit.

The inverse arrangement, where the constitutively expressed GFP utilises the host ribosome pool and the induced RFP utilises the o-ribosome pool (the o-RFP-h-GFP circuit), results in increased gene coupling with the isocost line gradient increasing by over six times in comparison to coupling in circuits using the host ribosomes (Figure 3c,d). Analysis of the model suggests that this is the result of competition for host ribosomal components, which are severely depleted. During translation of o-RFP, o-ribosomes are stabilised, which prevents release of ‘empty’ ribosomes, representing components of the ribosomal complex. This reduces the number of host ribosomes and so reduces host gene expression, including the expression of h-GFP (Figure S24).

We extended this approach of using multiple ribosome pools *in silico* by simulating the use of one o-ribosome pool for each circuit gene (Figure S25). Optimising the production of the two o-ribosome pools creates strong decoupling, with GFP falling less than 10% as RFP is induced. However, the RFP still shows a saturating response to increasing induction due to the finite o-ribosome pool size. There are also likely to be significant challenges in implementing multiple pools *in vivo* due to the cross talk found between different o-ribosome pools [15].

### Designing a dynamic resource allocation controller to uncouple co-expressed genes

Although we have shown that using separate host and o-ribosome pools can significantly reduce coupling between co-expressed genes, finite resource limitations still result in a saturated input-output response profile, while using a single orthogonal pool results in significant resource-mediated coupling. This coupling can, however, be exploited to design a feedback controller which acts to increase the circuit translational capability in line with increasing demand, thus alleviating *both* resource-mediated coupling and saturation effects (Figure 4). Increasing ribosome biosynthesis is not experimentally tractable so our controller acts to change the ratio of orthogonal to host ribosomes. We exploit the constitutive production of a repressor which utilises the oribosome pool for its own translation and inhibits the expression of the o-16S rRNA. Constitutive production of the repressor mRNA means that repressor protein levels act as a sensor for translational demand (Figure 4). To guide our design, we first implemented the feedback mechanism in our model (see Section S3.1.7). We demonstrate the function of our controller by considering the constitutive expression of one gene and simulating its response to the stepped induction of a second gene (Figure 4). When circuit demand is low, before the second gene induction, competition between the circuit and the controller is low (Figure 4b). This results in high expression of the controller and therefore high repression of the o-rRNA, meaning that few ribosomes are co-opted from the host. Upon induction of the second gene, the demand for o-ribosomes increases (Figure 4b). The repressor mRNAs will remain largely unaffected, but their translation falls due to increased competition (Figure 4b). The decrease in repressor production results in relief of the inhibition of the o-16S rRNA gene and so increased o-rRNA production and increased co-option of host ribosomes (Figure 4a). This results in the maintenance of circuit protein production as other circuit genes are induced (Figure 4c).

**Figure 4:**
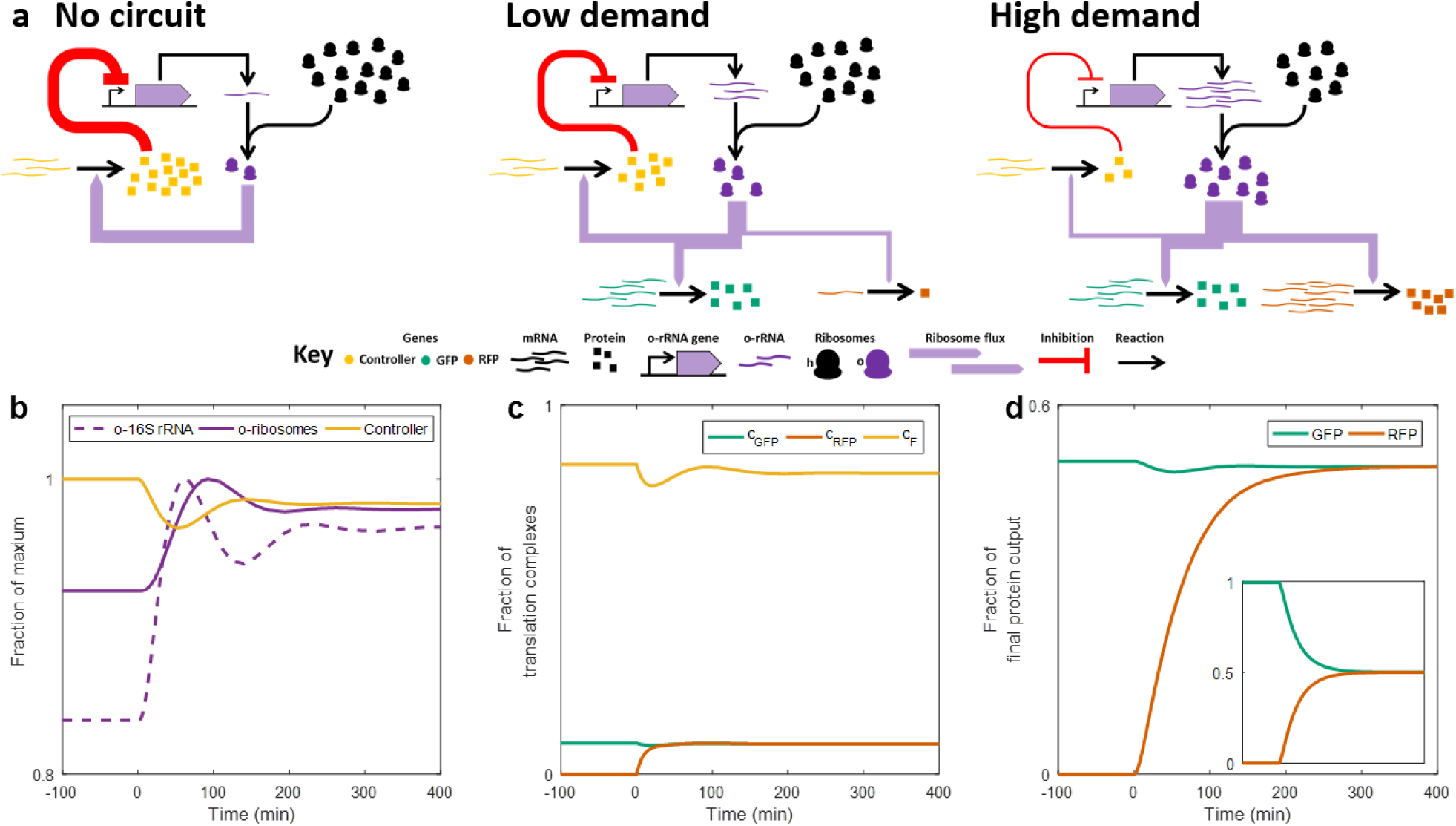
Operation of the negative feedback controller. *a.* Structure and function of the negative feedback controller. In the absence of the circuit genes, repression of o-rRNA production is high, and so co-option of ribosomes to the orthogonal pool is low. Upon the introduction of a low demand circuit the o-ribosome pool is redistributed to express both circuit and controller genes. This reduces translation of the constitutively expressed controller due to resource competition, and as a result o-rRNA transcription increases. As circuit demand increases (here RFP transcription increases), increased resource competition results in a greater fall in controller translation. This reduces the repression of the o-rRNA production, allowing more co-option of ribosomes from the host ribosome pool. **b.** Simulation showing the changing concentrations of the controller components in response to induction of RFP, which creates increased ribosomal demand. **c.** Changing distribution of the translation complexes, *c*, in response to RFP induction of genes utilising o-ribosomes for translation. *c_F_*, controller mRNA-o-ribosome translation complex. **d.** Protein output over time. Inset, coupling in the o-ribosome pool without controller. The induction rate of the o-rRNA was tuned so that both models produce the same protein concentration.

To test the robustness of our design to likely levels of uncertainty and variability arising from potential experimental implementations, we carried out sensitivity and robustness analyses around the optimal solution identified (Figure 5). This indicated that the feedback topology is highly robust, with all designs tested showing a better mRNA-protein mapping than the circuit using the o-ribosome pool without control, including those parameters which are difficult to design such as the *o_ρ_* (Figure 5a, inset). However, we do see the emergence of a trade-off between decoupling ability and gene expression, with decoupling coming at a cost to gene expression (Figure S26). Our sensitivity analysis shows that the controller needs to have a high ability to sequester o-ribosomes. Increasing the controller’s transcription rate and RBS strength is predicted to increase the ability of the controller to decouple genes (Figure 5b,c), with the former also raising gene expression while the latter reduces it. Varying the dissociation constant across a tenfold range allows the protein levels to be tuned at no cost to the level of decoupling achieved (Figure 5d), while varying the Hill function coefficient shows the need for the controller to be highly non-linear (Figure 5e).

**Figure 5:**
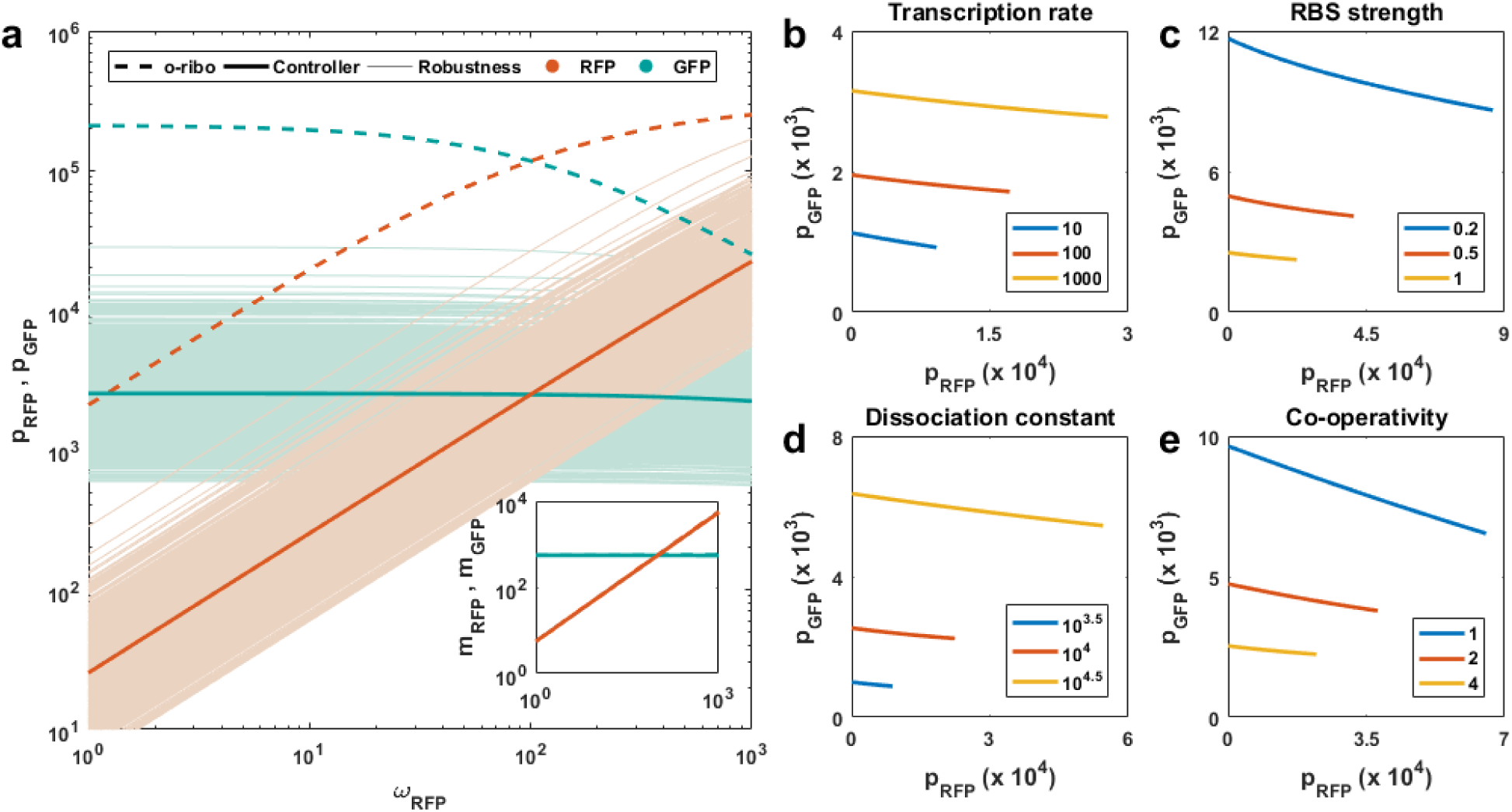
Design of the feedback controller. *a.* Function and robustness of the controller decoupling in comparison to using the o-ribosome pool alone. The optimal controller parameters were perturbed by drawing random values (*N* = 1000) between ±50% of the original value. All parameters controlling o-rRNA and controller protein were allowed to vary. Inset, mRNA levels of RFP and GFP. Legend explanation: o-ribo, uncontrolled o-ribosome pool (i.e. when *ω_F_* = 0 mRNAs per min); Controller, optimal feedback controller; Robustness, action of the perturbed controllers. **b.** Impact of varying the maximal transcription rate of the controller. **c.** Impact of varying the strength of the RBS **d.** Impact of varying the dissociation constant of the repressor. **e.** Impact of varying the Hill function coefficient of the repressor.

### Testing the controller *in vivo*

Having demonstrated the feasibility of using a feedback controller to dynamically allocate ribosomes between host and circuit, we implemented a prototype controller *in vivo*. Using the results of our sensitivity analysis as a guide, we based our controller on the strongly binding LacI repressor (*k_D_* ≈ 0.02 nM, [21]), which also shows a highly non-linear mode of action due to the dimerisation steps required to produce the functional complex. We selected the strong *P_LacI q_* promoter to drive LacI transcription [22] (Figure 5b), and used the same orthogonal RBS as previously. We deleted the host chromosomal LacI gene and placed the exogenous copy with promoter and o-RBS into the plasmid with the o-16S rRNA. Comparing the 0.5 mM IPTG treatment of the controlled circuit with the uncontrolled o-RFP-oGFP circuit of equivalent protein number (i.e. equivalent demand for ribosomes), we found that the controller decreases coupling by 50% (Figure 6). Tuning the controller threshold with ITPG allows the tuning of protein levels at no cost to decoupling (consistent with Figure 5d).

**Figure 6:**
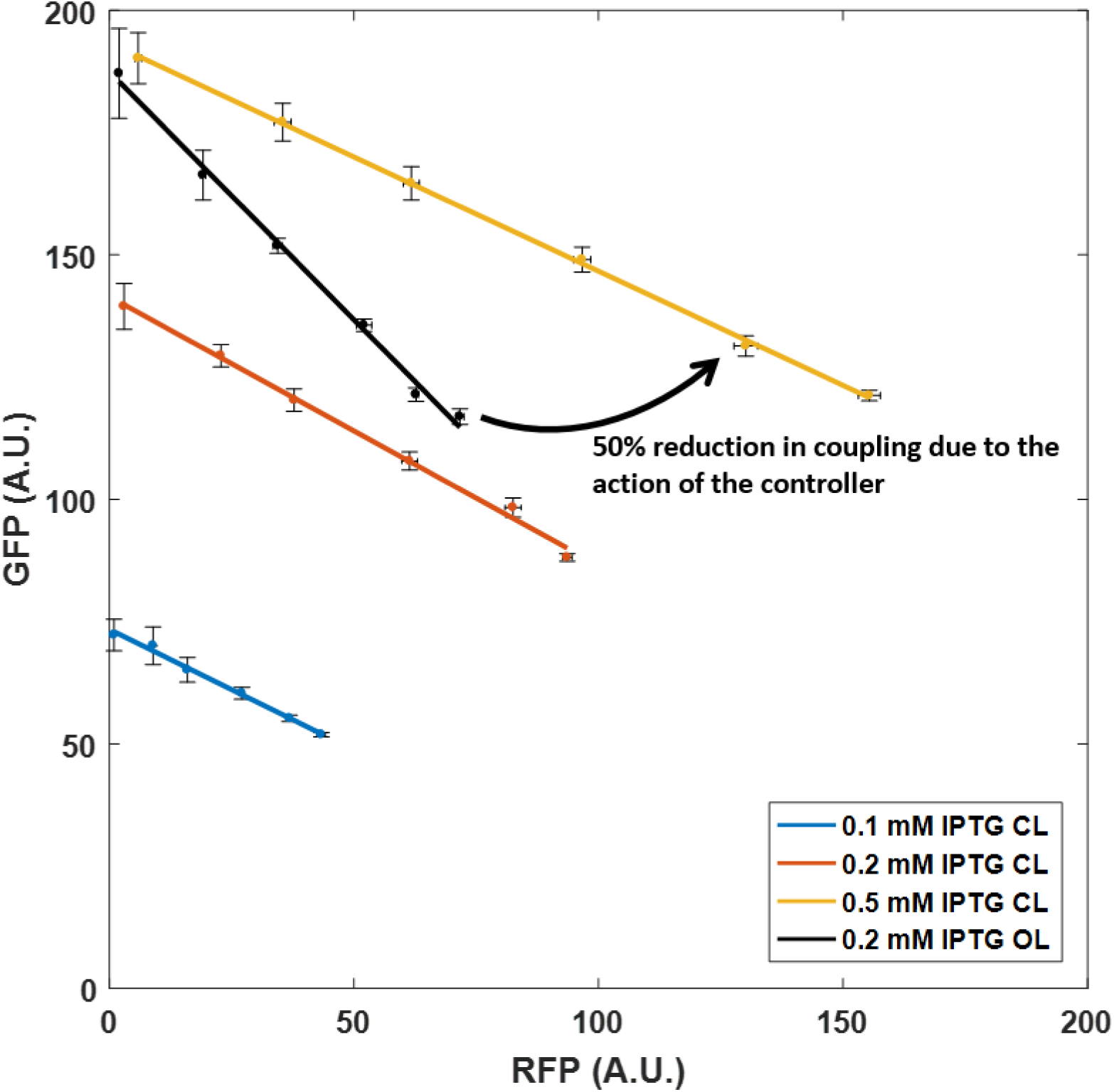
*In vivo* implementation of the prototype controller. Response of constitutively expressed o-GFP as o-RFP is induced with AHL. The action of the controller can be tuned with IPTG as shown (CL, closed loop). For comparison the o-RFP-o-GFP results which produce comparable protein levels are also shown (OL, open loop). Points are mean steady state fluorescence ±S.D. Isocost lines are fit to mean values.

## Discussion

To control cellular processes synthetic biologists and biotechnologists often use regulation of gene expression; by regulating transcription we assume protein levels will follow. However, the use of a common pool of cellular resources results in the emergence of hidden interactions, ‘couplings’, between genes which are not immediately apparent from circuit topologies. This can result in a breakdown in the relationship between transcriptional regulation (input) and protein levels (output). In this paper, we have shown the feasibility of dividing the cell’s translational resources to reverse this breakdown.

We have demonstrated that the use of orthogonal ribosomes is non-toxic to host cells (in agreement with previous reports), and that they can be used successfully for the targeted expression of synthetic circuit genes. Whilst we observe some crosstalk, with a small propensity of orthogonal ribosomes to translate host genes, when gene expression is low this does not appear to significantly impact on the design of our resource allocators and controller. Manipulating the size of the orthogonal ribosome pool provides an additional ‘dial’ for controlling gene expression [23].

By carefully selecting which genetic modules utilise either the host or orthogonal ribosome pools we can reduce hidden interactions between co-expressed genes (Figure S18A). Our simple model of cellular physiology is sufficient to determine which modules should utilise the o-ribosome pool to achieve the best coupling reduction. The greatest level of decoupling is achieved when constitutively expressed genes utilise the orthogonal pool while dynamically regulated genes utilise the host pool. In this manner, the orthogonal pool appears to act as an insulation device.

However, this use of multiple ribosome pools does not prevent resource-mediated saturation. The response of a genetic module to increasing input does not show increasing output due to lack of additional free ribosomes at high levels of input, which results in the saturating response and effective autoinhibition of each gene. Taking inspiration from control theory, we took advantage of gene couplings to design a negative feedback controller which acts to dynamically divert resources to the synthetic circuit as demand for them increases. This controller acts to remove both resource mediated saturation and gene-gene coupling. Guided by the analysis of our model, we implemented a prototype controller, based on the repressor lacI, and found that it could reduce gene-gene coupling in a simple two gene circuit by 50%. The decoupling ability of the controller is robust, with only protein levels, not coupling, being changed with the addition of IPTG. Although our model predicts that coupling can be reduced further than the 50% observed in our prototype design, the remaining uncontrolled coupling observed experimentally is likely to be due to factors not explicitly accounted for in our model, such as RNA polymerases, or specific tRNAs and/or amino acids.

In previous studies, a number of *transcriptional* resource allocators have been developed, either by directing the core polymerase via use of orthogonal *σ* factors (analogous to our use of multiple ribosome pools), or by using feedback to control RNA polymerase expression across species [24, 25]. In another approach, Venturelli *et al.* proposed and implemented a global resource allocator based on increasing decay rates of host mRNAs to reduce ribosomal competition [26]. Our dynamic resource allocator complements these efforts by providing another potential layer of control at the translational level, which multiple studies have shown is the key limiting factor in determining bacterial gene expression. Our results show that, by using the power of feedback control, translational capacity can be controlled in order to optimally manage the consumption of host resources and reduce circuit context dependency. These decoupling mechanisms provide designers with multiple means to reduce host-circuit interactions and improve circuit modularity, without the need for costly and time-consuming rounds of experimentation and library screening, thus facilitating the real-world implementation of synthetic gene circuits in future biotechnological applications.

## Acknowledgements

The authors are indebted to Prof. Jason Chin (MRC-Laboratory of Molecular Biology) and Dr Amit Sachdeva (University of East Anglia) for the plasmids encoding the orthogonal 16S RNA. The authors also thank Dr Domitilla Del Vecchio (Massachusetts Institute of Technology) and Prof. Victor de Lorenzo (Centro Nacional de Biotechnoloǵıa, CSIC) for the plasmid MBP1.0 and strain *E. coli* DH5*α λ pir* respectively. APSD and DGB acknowledge funding from the University of Warwick and the EPSRC & BBSRC Centre for Doctoral Training in Synthetic Biology (grant EP/L016494/1). JK and JIJ acknowledge support from the European Union’s Horizon 2020 research and innovation programme (grant no. 633962) and support from the Biotechnology and Biological Sciences Research Council (BBSRC) (grant BB/M009769/1).

## Methods

### Mathematical modelling

The full model and modifications are described in detail in Section S3. All ordinary differential equation models were simulated to steady state using custom software written in MATLAB 2016a (The MathWorks Inc., MA, USA).

### Cloning procedures and construction of reporter strains

The characteristics of the bacteria, plasmids, and primers used in this study are described in Figure S1, Table S1 and Table S2. DNA manipulation was carried out following standard protocols [27]. Plasmid DNA was isolated from bacterial cells using a commercial QIAprep Spin Miniprep Kit (Qiagen, Hilden, Germany). Restriction endonucleases were purchased from New England Biolabs (NEB, Ipswich, MA, USA).

Plasmid pSEVA63-Dual was derived from MBP 1.0, the original circuit containing the genes for constitutive GFP and inducible RFP expression [6]. The circuit was amplified using the pair of primers Dual F/-R, digested with PacI/SacI and cloned into pSEVA631 using the same restriction endonucleases [28].

Two subsequent cloning steps were carried out to replace the original RBS sequence of RFP in the plasmid. First, the original RBS of RFP was replaced with the orthogonal sequence (ACAATTTTCATATCCCTC-CGCAA) using the Fast Cloning method [29] with primers o-RFP F/-R resulting in the plasmid pEMG-o-RFP-h-GFP. The RBS of GFP was replaced in this circuit using the pair of primers Dual F and Dual R and the PCR product was cloned into pSEVA631 as described above, which resulted in the plasmid pSEVA63-o-RFP-h-GFP. Plasmids pSEVA63-h-RFP-o-GFP and pSEVA63-o-RFP-o-GFP were generated with a similar approach in this case using primers o-GFP F/-R.

The RBS of the *lacI* gene present in the plasmid pRSF ribo-Q1 O-*gst-cam* [20] was replaced with the orthogonal version also following this strategy: three partial fragments representing the whole plasmid were amplified by PCR with the respective primer pairs (pAS2 o-lacI pt F/-R, pAS2 o-lacI pt2 F/-R, pAS2 o-lacI pt3 F/-R and pAS2 o-lacI pt4 F/-R). Each product had overlapping regions and one of these regions contained the orthogonal RBS directly upstream of the *lacI* gene. DpnI treatment was carried out to remove the template DNA after the PCR reaction, and the DNA fragments were ligated by isothermal assembly [30] yielding plasmid pRSF ribo-Q1 o-gst-cam o-lacI.

The MG1655ΔlacIZYA strain – this is the parental MG1655 strain lacking the *lacI* gene and the *lac* operon (i.e. the *lacZYA* genes) – was constructed by replacing the target genomic regions with a kanamycin antibiotic cassette (pKD4) using primers lac KO F/-R, followed by removal of the kanamycin resistance using the FLP recombinase using a previously described method [31, 32]. Genomic deletions were confirmed by PCR and by ensuring that the activity of *β*-galactosidase was not present in the strain.

All plasmid constructs described above were introduced into either *E. coli* DH5*α* or DH5*αλpir* strains by transformation for DNA amplification. All experiments in this paper were performed in *E. coli* MG1655 after transformation with the corresponding plasmids with the exception of those involving the feedback controller (pRSF ribo-Q1 o-gst-cam o-lacI), which were carried out in MG1655ΔlacIZYA to prevent cross talk between the endogenous *lacI* gene and the controller.

### Cell culturing and fluorescence measurements

*E.coli* strains used in this study were always grown in Luria-Bertani (LB) medium at 37°C. The antibiotics kanamycin (50 *µ*g mL^−1^), ampicillin (150 *µ*g mL^−1^) and gentamicin (20 *µ*g mL^−1^) were added when necessary. In a typical experiment, for each of the biological replicates a single colony from the strain of interest was taken from a fresh plate and cultured overnight in

1 mL of liquid medium in 24-well plates (500 rpm, PMS-1000i Microplate Shaker, Grant, Shepreth, UK). These cultures were diluted 500-fold in the same medium and further grown in the same conditions. Once they reached the exponential phase (OD600 = 0.2 − 0.3), 2 *µ*L of the culture were transferred into 1 mL of fresh medium containing Isopropyl *β*-D-1-thiogalactopyranoside (IPTG, Sigma-Aldrich, St. Louis, MO, USA, final concentrations of 0.1, 0.2 and 0.5 mM) and/or N-acyl homoserine lactone (AHL, Sigma-Aldrich, St. Louis, MO, USA, final concentrations of 1.25, 2.5, 5, 10, 20 nM) and cultured in the same way as in the previous two steps.

Growth and fluorescence of these cultures was monitored over time as follows: every hour the OD_600_ of the cultures was determined using a CLARIOstar microplate reader (BMG Labtech, Ortenberg, Germany). At the same time, 50 *µ*L aliquots were taken from each well mixed with 150 *µ*L of PBS. The volume of the culture in the wells was kept constant by replenishing with the same volume of fresh medium including the same concentration of the inducers. The suspension of cells in PBS was loaded into an Attune NxT Flow Cytometer (ThermoFisher, Waltham, MA, USA) and analysed for GFP and RFP expression using blue (excitation 561 nm; emission 620/15 nm) and yellow (excitation 561 nm; emission 620/15 nm) lasers respectively. For each sample, 20,000 events were analysed and populations means were estimated using the default software of the instrument.

## References

1. J. A. Brophy and C. A. Voigt, “Principles of genetic circuit design.,” Nature Methods, vol. 11, no. 5, pp. 508–20, 2014.

2. S. Cardinale and A. P. Arkin, “Contextualizing context for synthetic biology - identifying causes of failure of synthetic biological systems,” Biotechnology Journal, vol. 7, no. 7, pp. 856–866, 2012.

3. Y. Qian, H.-H. Huang, J. Jiménez, and D. Del Vecchio, “Resource competition shapes the response of genetic circuits,” ACS Synthetic Biology, 2017.

4. M. Scott, C. W. Gunderson, E. M. Mateescu, Z. Zhang, and T. Hwa, “Interdependence of cell growth and gene expression: origins and consequences.,” Science, vol. 330, no. 6007, pp. 1099–1102, 2010.

5. F. Ceroni, R. Algar, G.-B. Stan, and T. Ellis, “Quantifying cellular capacity identifies gene expression designs with reduced burden,” Nature Methods, vol. 12, no. 5, pp. 415–423, 2015.

6. A. Gyorgy, J. I. Jim´enez, J. Yazbek, H.-H. Huang, H. Chung, R. Weiss, and D. Del Vecchio, “Isocost Lines Describe the Cellular Economy of Genetic Circuits,” Biophysical Journal, vol. 109, no. 3, pp. 639–646, 2015.

7. M. Carbonell-Ballestero, E. Garcia-Ramallo, R. Montanez, C. Rodriguez-Caso, and J. Marcia, “Dealing with the genetic load in bacterial synthetic biology circuits: convergences with the Ohm’s law,” Nucleic Acids Research, pp. 1–12, 2015.

8. T. E. Gorochowski, I. Avcilar-Kucukgoze, R. A. Bovenberg, J. A. Roubos, and Z. Ignatova, “A minimal model of ribosome allocation dynamics captures trade-offs in expression between endogenous and synthetic genes,” ACS Synthetic Biology, vol. 5, pp. 710–720, 2016.

9. A. Hamadeh and D. del Vecchio, “Mitigation of resource competition in synthetic genetic circuits through feedback regulation,” Proc. of IEEE Conference on Decision and Control, 2015.

10. Y. Qian and D. Del Vecchio, “Mitigation of ribosome competition through distributed sRNA feedback,” Proc. of IEEE Conference on Decision and Control, 2016.

11. T. Shopera, L. He, T. Oyetunde, Y. J. Tang, and T. S. Moon, “Decoupling resource-coupled gene expression in living cells,” ACS Synthetic Biology, 2017.

12. C. Tan, P. Marguet, and L. You, “Emergent bistability by a growth-modulating positive feedback circuit,” Nature Chemical Biology, vol. 5, pp. 842–8, nov 2009.

13. D. E. Cameron and J. J. Collins, “Tunable protein degradation in bacteria.,” Nature Biotechnology, vol. 32, no. 12, pp. 1276–1281, 2014.

14. A. Hui and H. de Boer, “Specialized ribosome system: preferential translation of a single mRNA species by a subpopulation of mutated ribosomes in Escherichia coli.,” Proceedings of the National Academy of Sciences of the United States of America, vol. 84, no. 14, pp. 4762–6, 1987.

15. O. Rackham and J. W. Chin, “A network of orthogonal ribosome*·*mRNA pairs,” Nature Chemical Biology, vol. 1, no. 3, pp. 159–166, 2005.

16. L. M. Chubiz and C. V. Rao, “Computational design of orthogonal ribosomes,” Nucleic Acids Research, vol. 36, no. 12, pp. 4038–4046, 2008.

17. A. Espah Borujeni, A. S. Channarasappa, and H. M. Salis, “Translation rate is controlled by coupled trade-offs between site accessibility, selective RNA unfolding and sliding at upstream standby sites,” Nucleic Acids Research, vol. 42, no. 4, pp. 2646–2659, 2014.

18. B. J. Des Soye, J. R. Patel, F. J. Isaacs, and M. C. Jewett, “Repurposing the translation apparatus for synthetic biology,” Current Opinion in Chemical Biology, vol. 28, pp. 83–90, 2015.

19. A. Y. Weiße, D. A. Oyarzún, V. Danos, and P. S. Swain, “Mechanistic links between cellular trade-offs, gene expression, and growth,” Proceedings of the National Academy of Sciences, vol. 112, no. 9, pp. E1038–E1047, 2015.

20. A. Sachdeva, K. Wang, T. Elliott, and J. W. Chin, “Concerted, rapid, quantitative, and site-specific dual labeling of proteins,” Journal of the American Chemical Society, vol. 136, no. 22, pp. 7785–7788, 2014.

21. C. M. Falcon and K. S. Matthews, “Operator DNA sequence variation enhances high affinity binding by hinge helix mutants of lactose repressor protein,” Biochemistry, vol. 39, no. 36, pp. 11074–11083, 2000.

22. P. Penumetcha, K. Lau, X. Zhu, K. Davis, T. T. Eckdahl, and a. M. Campbell, “Improving the Lac System for Synthetic Biology,” Bios, vol. 81, no. 1, pp. 7–15, 2010.

23. J. A. J. Arpino, E. J. Hancock, J. Anderson, M. Barahona, G. B. V. Stan, A. Papachristodoulou, and K. Polizzi, “Tuning the dials of synthetic biology,” Microbiology, vol. 159, no. 7, pp. 1236–1253, 2013.

24. T. H. Segall-Shapiro, A. J. Meyer, A. D. Ellington, E. D. Sontag, and C. A. Voigt, “A ‘resource allocator’ for transcription based on a highly fragmented T7 RNA polymerase.,” Molecular Systems Biology, vol. 10, no. 7, p. 742, 2014.

25. M. Kushwaha and H. M. Salis, “A portable expression resource for engineering cross-species genetic circuits and pathways,” Nature Communications, vol. 6, p. 7832, 2015.

26. O. S. Venturelli, M. Tei, S. Bauer, L. J. G. Chan, C. J. Petzold, and A. P. Arkin, “Programming mRNA decay to modulate synthetic circuit resource allocation,” Nature Communications, vol. 8, pp. 1–26, 2017.

27. J. Sambrook and D. Russel, Molecular Cloning: A Laboratory Manual. Cold Spring Harbor Laboratory Press, 2001.

28. R. Silva-Rocha, E. Martínez-García, B. Calles, M. Chavarría, A. Arce-Rodríguez, A. de Las Heras, A. D. Páez-Espino, G. Durante-Rodríguez, J. Kim, P. I. Nikel, R. Platero, and V. de Lorenzo, “The Standard European Vector Architecture (SEVA): a coherent platform for the analysis and deployment of complex prokaryotic phenotypes.,” Nucleic Acids Research, vol. 41, no. Database issue, pp. D666–75, 2013.

29. C. Li, A. Wen, B. Shen, J. Lu, Y. Huang, and Y. Chang, “FastCloning: a highly simplified, purification-free, sequence- and ligation-independent PCR cloning method,” BMC Biotechnology, vol. 11, p. 92, oct 2011.

30. D. G. Gibson, L. Young, R.-Y. Chuang, J. C. Venter, C. A. Hutchison, and H. O. Smith, “Enzymatic assembly of DNA molecules up to several hundred kilobases,” Nature Methods, vol. 6, pp. 343–345, may 2009.

31. K. Datsenko and B. Wanner, “One-step inactivation of chromosomal genes in Escherichia coli K-12 using PCR products,” Proceedings of the National Academy of Sciences USA, vol. 97, no. 12, pp. 6640–6645, 2000.

32. T. Baba, T. Ara, M. Hasegawa, Y. Takai, Y. Okumura, M. Baba, K. A. Datsenko, M. Tomita, B. L. Wanner, and H. Mori, “Construction of Escherichia coli K-12 in-frame, single-gene knockout mutants: the Keio collection,” Molecular Systems Biology, vol. 2, feb 2006.

